# The ‘near T2T’ Canton-S assembly is inconsistent with other Canton-S genomes

**DOI:** 10.64898/2026.11.19.767897

**Authors:** J.J. Emerson

## Abstract

Canton-S is among the most widely used *Drosophila melanogaster* strains, underlying substantial community infrastructure, including serving as a wild-type strain for: genetics studies^1^; studying phenotypes^2–4^, especially in behavioral studies^5–7^; and complex trait mapping studies^8^ among many others. The recent ‘near T2T’ assembly of Canton-S (LIU et al. ^9^) is presented as delivering modern sequencing and assembly quality to the *Drosophila* community via this important strain. I find that this assembly is inconsistent with three other independently sequenced Canton-S genome assemblies on a magnitude comparable to differences between distinct strains, raising concerns about its genetic identity that the community should consider when choosing whether to adopt it as a resource.

## Introduction

The recently published study of *D. melanogaster* by LIU et al.^9^ claims to deliver a nearly complete genome assembly of the widely used strain Canton-S, exceeding that of the current reference assembly of the *Drosophila* community, the Release 6 assembly of strain iso-1^10^, and in the process, closing many gaps. Interestingly, this important strain had been previously assembled to a standard comparable to Release 6 at least twice before: CHAKRABORTY et al.^11^ and WIERZBICKI et al.^12^. More recently, the Drosophila Laboratory Pangenome Database (DLPD) released a modern assembly of Canton-S, with data available in July 2024^13^. Genetic background modulates phenotype, so a community resource is only as useful as the certainty with which its genotype can be tied to its name. It is well understood that variation among independently maintained stocks may lead to different conclusions among studies that fail to employ genetically identical strains^14,15^. Such variation may arise from stock contamination (i.e., substitution or admixture with a different *D. melanogaster* strain), a persistent risk for lab-maintained fly stocks that motivates authentication of key strains. The existence of four independent assemblies of Canton-S allowed me to quantify the differences between strains obtained at different times and maintained in different laboratories. These assemblies have four independent origins from the same Bloomington stock center source: by no later than 2006 (A.D. Long, personal communication), the Drosophila Synthetic Population Resource (DSPR) incorporated Canton-S from the stock center to serve as one of its 15 founder lines (line A1)^8^, which was later sequenced and assembled by CHAKRABORTY et al.^11^; in January 2018 (R. Kofler, personal communication), WIERZBICKI et al.^12^ sourced Canton-S from Bloomington in order to study piRNA clusters; in November, 2023, Mahul Chakraborty obtained Canton-S from Bloomington for the DLPD project (personal communication); and in 2018, the stock was ordered from Bloomington by the laboratory of Dr. Yi Zhong, from whom LIU et al.^9^ obtained it for genome sequencing and assembly. Notably, Bloomington’s Canton-S stock was found to be contaminated in 2016, prompting reacquisition from the Kyoto Drosophila Stock Center, which held a copy of the original Bloomington stock^16^. CHAKRABORTY’s pre-2016 stock therefore represents Bloomington’s original Canton-S, while the remaining three assemblies derive from Kyoto’s copy, making this comparison capable of quantifying any genetic variation accumulated between the two independently maintained stocks.

## Methods

### Inter-assembly distance calculations and visualization of assembly relationships

To measure similarity between assemblies, I compared the k-mer inventories of their euchromatic sequence. Euchromatin boundaries defined on the iso-1 Release 6 reference^10^ were lifted over to the 20 other assemblies (minimap2 and paftools.js), and the homologous euchromatic segments were extracted. Canonical 21-mers in each assembly’s euchromatic sequence were counted with Jellyfish, and those occurring exactly once (unique k-mers) were retained. For all 210 pairwise comparisons among the 21 assemblies, the Jaccard similarity of the two unique k-mer sets was calculated, and from it the Mash distance D^17^ as an estimate of pairwise nucleotide diversity. The full assembly × assembly matrix of Mash distances was used to construct a neighbor-joining tree with ape (nj) in R, and the resulting unrooted tree was plotted with ggtree.

### Chromosome-wide variation analysis

To assess the structure of variation directly, I constructed pairwise alignments for a curated set of assembly pairs. The set includes positive and negative controls for the expected level of variation between two *Drosophila* strains: assemblies sharing a strain label are predicted to be virtually devoid of variation, while those with different labels should show variation throughout the genome. Where segments identical by descent (IBD; i.e., sharing recent common ancestry) are interspersed with non-IBD segments, the prediction is a block-like mixture of tracts with typical *Drosophila* variation (∼0.5%) and tracts with none (∼0%).

The euchromatic segments extracted for the k-mer analysis were aligned with minimap2 (asm10 preset), and single-nucleotide substitutions were called with paftools.js call in regions covered by exactly one alignment. Variation was summarized as substitutions per callable base in 100 kb windows with a 10 kb slide and plotted with matplotlib in Python.

## Results & Discussion

I estimated pairwise nucleotide diversity 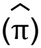 via Mash distance^17^ across 21 *D. melanogaster* assemblies, including the four Canton-S assemblies above (210 pairwise comparisons in total) using singleton 21-mers from the five major Muller element scaffolds (i.e., the major chromosome arms: X, 2L, 2R, 3L, and 3R). Despite being sourced independently from the stock center in 2006, 2018, and 2023, CHAKRABORTY’s^11^, WIERZBICKI’s^12^, and DLPD’s^13^ Canton-S assemblies have nearly identical unique k-mer inventories, leading to an extremely low estimate of pairwise nucleotide diversity 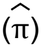 (Fig. 1a). On the other hand, LIU’s^9^ Canton-S assembly diverges from the other three assemblies by nearly as much as unrelated strains, far exceeding intra-strain variation observed elsewhere in the panel (Fig. 1a). In a neighbor-joining tree of the pairwise 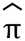 estimates above, LIU’s assembly is a sister to the nearly identical CHAKRABORTY/WIERZBICKI/DLPD trio (Fig. 1b), suggesting not only that the LIU strain is closely related to the other Canton-S strains, but also exhibits substantial differences unexpected when comparing two independent assemblies of the same strain (Fig. 1a).

**Figure 1.**
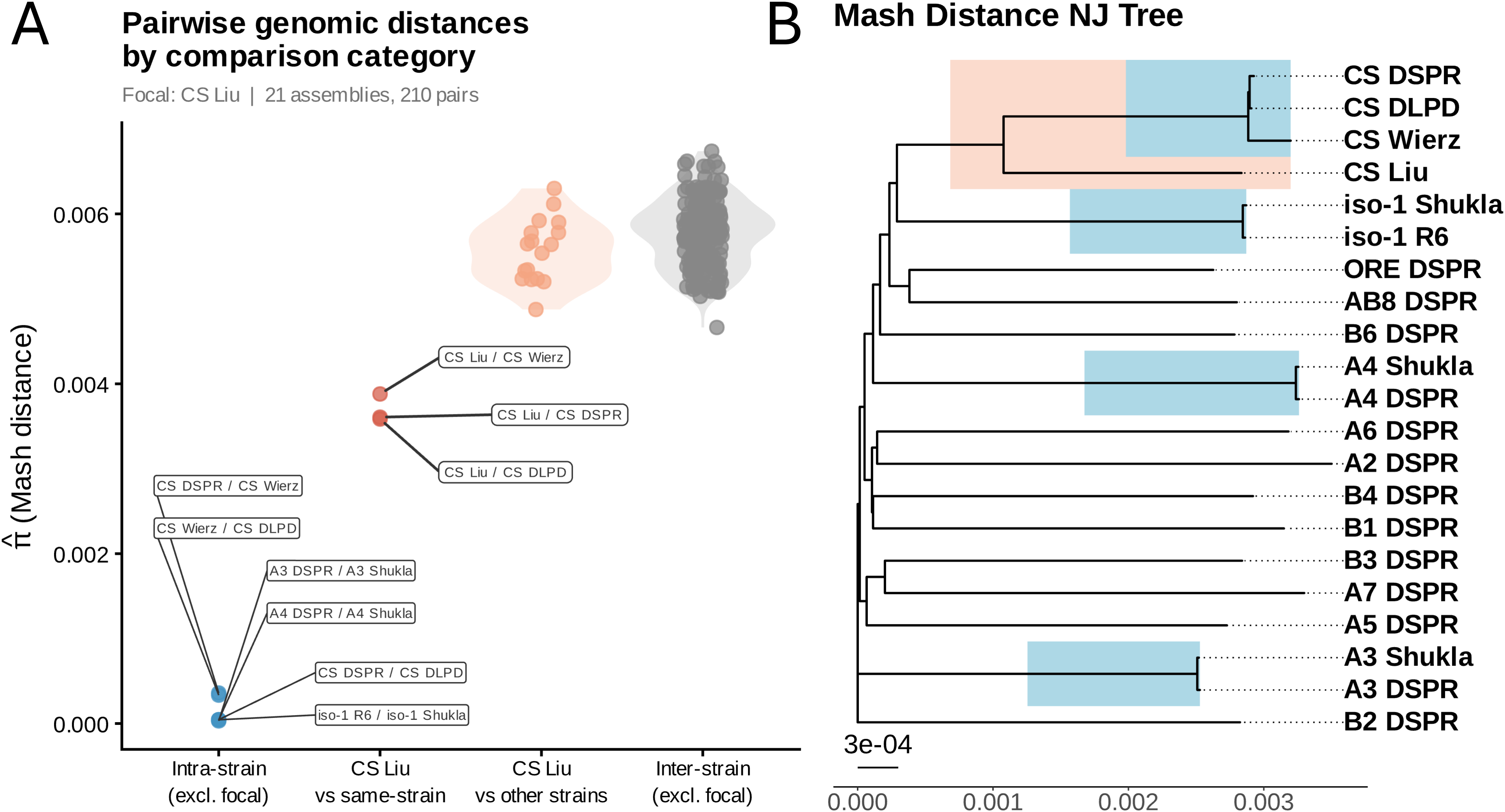
(A) Swarm plot of all pairwise Mash distances, grouped into four categories: within-strain-label assembly comparisons excluding Canton-S from Liu et al. 2026; Canton-S from Liu et al. 2026 vs. other Canton-S assemblies; Canton-S from Liu et al. 2026 vs. non-Canton-S assemblies; and all remaining between-strain-label assembly comparisons. Axis labels abbreviate these as “Intra-strain (excl. focal)”, “CS Liu vs same-strain”, “CS Liu vs other strains” and “Inter-strain (excl. focal)”, where focal denotes CS Liu. (B) Neighbor-joining tree built from the same all-by-all Mash D distance matrix. Blue boxes mark clades possessing the same strain label and nearly no divergence; the peach box marks a clade containing all assemblies labeled Canton-S. Tip labels combine each assembly’s strain designation with an abbreviation for its source: DSPR = Chakraborty et al. 2019; Wierz = Wierzbicki et al. 2022; DLPD = Drosophila Laboratory Pangenome Database (unpublished); Shukla = Shukla et al. 2025; R6 = Hoskins et al. 2015 (Release 6 reference); Liu = Liu et al. 2026.

Windowed π across euchromatic Muller elements shows the variable sites distinguishing LIU’s assembly from the other three are block-structured, rather than randomly distributed (Fig. 2), alternating between discrete tracts of high nucleotide diversity comparable to levels found between different strains and near-zero tracts, consistent with discrete haplotype blocks rather than uniform accumulated variation. Additionally, given biologically reasonable mutation rates, mutation accumulation alone is incapable of introducing enough variation to be mistaken for inter-strain variation over even hundreds of generations.

**Figure 2.**
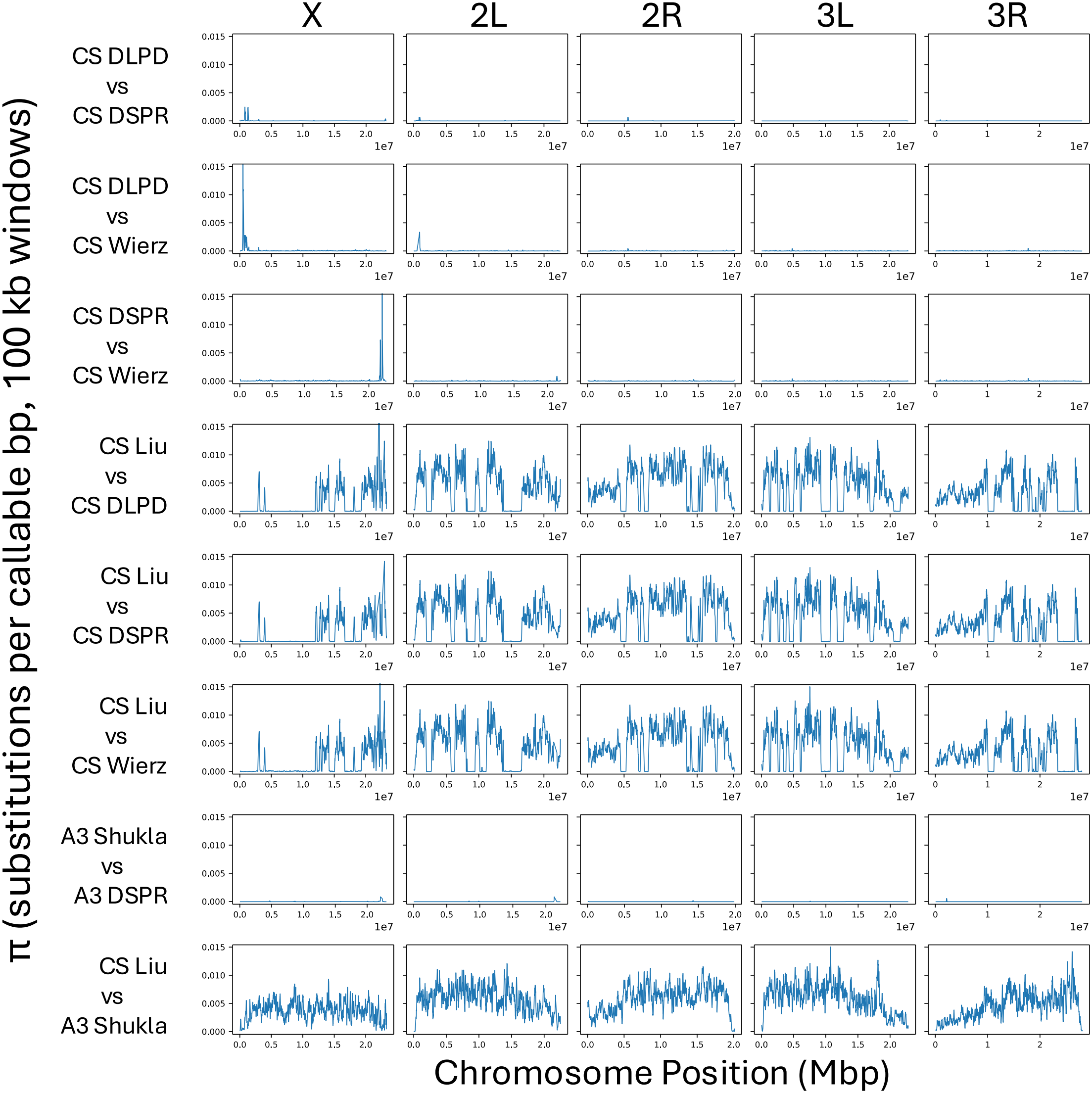
Alignment-based windowed nucleotide diversity (π) across euchromatic Muller elements, comparing pairs of independently sourced genome assemblies. Rows 1–3 establish an intra-Canton-S baseline excluding the focal Canton-S assembly by Liu et al. 2026 (DLPD, unpublished; Chakraborty et al. 2019; Wierzbicki et al. 2022). Rows 4–6 compare the focal assembly (Liu et al. 2026) against each of those same three non-focal Canton-S assemblies. Row 7 is an intra-strain control for a second strain, A3: Chakraborty et al. 2019 vs. Shukla et al. 2025. Row 8 is an inter-strain control: Liu et al. 2026 (Canton-S) vs. Shukla et al. 2025 (A3). All eight rows share the same y-axis scale (0–0.015) for direct visual comparison. The x-axis is the position relative to the euchromatin extracted from the assembly named first in the comparison. Note that y-axis values above 0.015 are truncated.

These results are consistent with strain contamination introduced into the LIU stock at some point preceding its sequencing, though my approach cannot determine whether this would have occurred before or after the stock was procured from the stock center. One could also invoke substantial residual heterozygosity segregating in the stock center strain and captured by only one of the four independent assemblies, but this explanation is considerably less parsimonious. Given that CHAKRABORTY, WIERZBICKI, and DLPD are nearly identical and LIU is substantially different, residual heterozygosity is implausible, especially since one might predict under that model that LIU and WIERZBICKI would cluster together given how close in time they were sampled. Regardless of the cause of the differences, users should not assume genomic identity between the LIU assembly and other genomes carrying the Canton-S designation. Perhaps equally importantly, Bloomington’s current Canton-S stock, despite having been re-derived from Kyoto’s independently-held copy following the 2016 contamination event (Figure S1), is nearly identical to a Bloomington-sourced Canton-S sample from 2006 (Fig. 1a; Fig. 2, rows 1–3), predating that detour entirely. This indicates the original lineage has remained genetically stable across nearly two decades, including an extended period maintained at a separate institution.

The overarching goal LIU et al. set out to achieve, a modern, high-quality assembly of a foundational community strain, is a valuable one. While I cannot conclude that LIU achieved this goal, this does not leave the field without a modern assembly of a community strain: SHUKLA et al.^18^ already delivered three other assemblies employing modern long-read methodology, including the community reference strain iso-1, with contiguity and completeness comparable to LIU’s own. For Canton-S specifically, however, CHAKRABORTY et al.’s 2019 assembly^11^ and WIERZBICKI et al.’s 2022 assembly^12^ remain, to my knowledge, the best published representations of the strain while the DLPD represents an unpublished version of the strain employing current state of the art sequencing and assembly approaches.

I recommend that the *Drosophila* genetics and genomics communities treat the results shared by LIU as of uncertain provenance. Furthermore, given that the LIU assembly is mosaic rather than a pure representation of Canton-S, and that pure assemblies already exist, I recommend against using the LIU assembly for studies where strain provenance is critical, particularly when a Canton-S strain is desired. Finally, I recommend that assemblies claiming to represent strains widely used by the community, especially important strains like Canton-S, include genomic authentication as a condition of making that claim, consistent with established policies on authentication of key biological resources adopted by some funding agencies^19^. Strain designations alone do not guarantee genotype identity across independently maintained lab stocks, and every study that relies on an unverified strain inherits that uncertainty.

## Contributions

J.J.E. conceived the study, conceived and prototyped each analysis, and wrote the paper. J.J.E. supervised Claude Code in refactoring the analysis prototypes, generating the orchestration scripts, and constructing the GitHub repository for the sharing of methods and code (https://github.com/jjemerson/canton-s-authentication). J.J.E. is solely responsible for the content of the repository.

## Code availability

All analysis code, configuration files and pipeline documentation are available at https://github.com/jjemerson/canton-s-authentication. The version used in this study (v1.0.1) is archived at Zenodo^20^.

## Acknowledgements

J.J.E. acknowledges H.G. Shukla, M. Chakraborty, E. Cryan, and Andy Kern for comments on a draft of the manuscript and A.D. Long, R. Kofler, and M. Chakraborty for confirming the date and source of their Canton-S strains and M. Chakraborty for sharing his Canton-S assembly before publication.

