## Supplementary Figure S1 for "The ‘near T2T’ Canton-S assembly is inconsistent with other Canton-S genomes"

Provenance of stocks labeled Canton-S across stock centers and laboratories. Time runs left to right from 2002 onward; the 1987–2002 interval is compressed, marked by a break in the Bloomington line. Vertical arrows are stock transfers, colored by the lane they leave from; horizontal lines are propagation within a lane. Circles at the beginning of an arrow mark an institution or lab sharing a stock while those at the end of an arrow mark an institution or lab taking custody of a stock; squares mark public deposition of a genome assembly from that lineage (NCBI deposition dates; DLPD from its GitHub repository). The dotted stretch on the Bloomington line marks the contamination of its resident stock, whose onset is undocumented, resolved in 2016 by a transfer back from Kyoto's independently maintained copy.

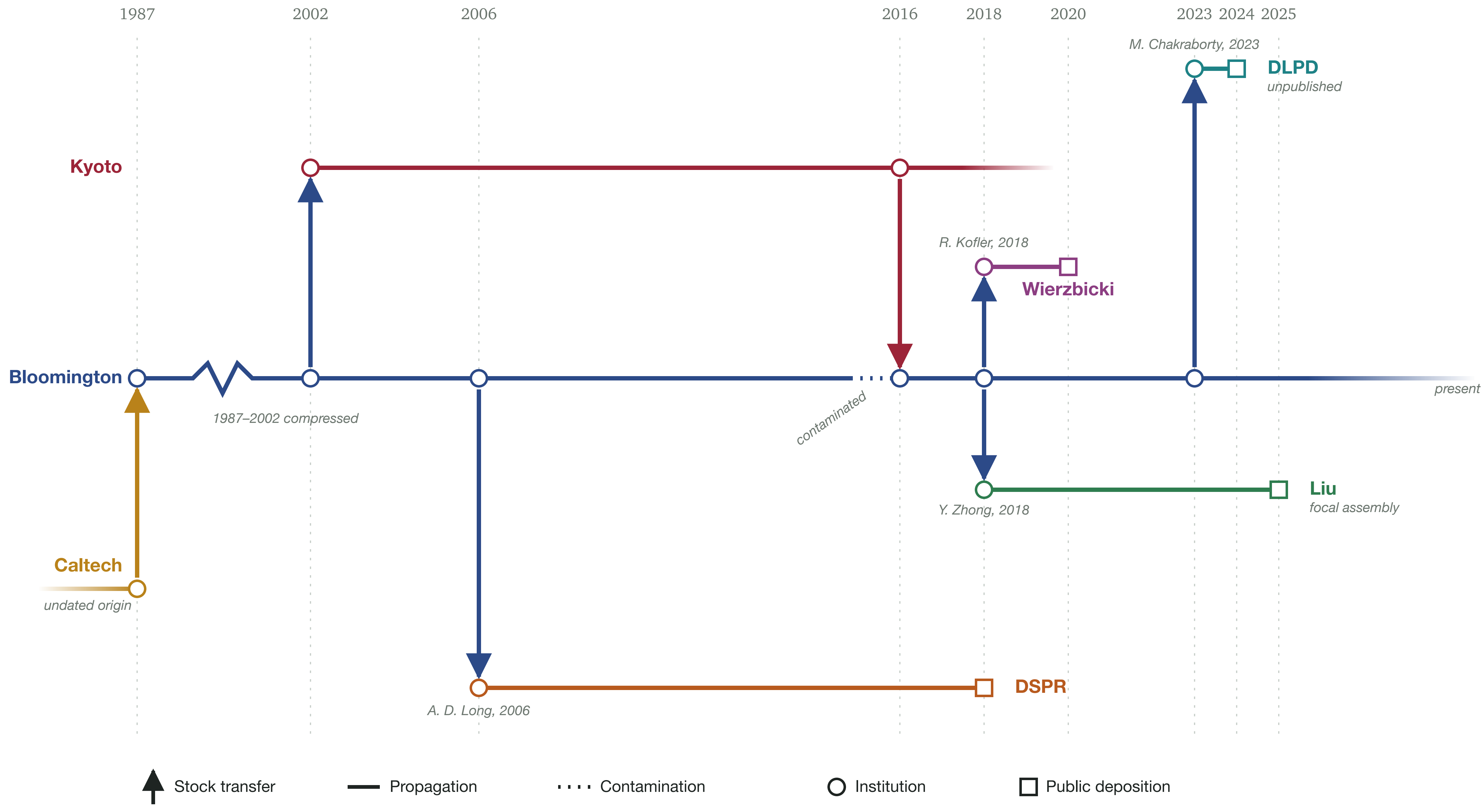
1987
2002
2006
2016
2018
2020
2023
2024
2025
M. Chakraborty, 2023
DLPD
unpublished
Kyoto
R. Kofler, 2018
Wierzbicki
Bloomington
1987–2002 compressed
contaminated
present
Liu
focal assembly
Y. Zhong, 2018
Caltech
undated origin
A. D. Long, 2006
DSPR
Stock transfer
Propagation
Contamination
Institution
Public deposition
